# PathFinder: a novel graph transformer model to infer multi-cell intra- and inter-cellular signaling pathways and communications

**DOI:** 10.1101/2024.01.13.575534

**Authors:** Jiarui Feng, Michael Province, Guangfu Li, Philip R.O. Payne, Yixin Chen, Fuhai Li

**Affiliations:** Institute for Informatics (I2), Washington University School of Medicine, Washington University in St. Louis, St. Louis, MO, USA; Department of Computer Science and Engineering, Washington University in St. Louis, St. Louis, MO, USA; Division of Statistical Genomics, Department of Genetics, Washington University in St. Louis, St. Louis, MO, USA; Department of Surgery, University of Missouri-Columbia, Columbia, MO, 65212, USA; Department of Molecular Microbiology and Immunology, University of Missouri-Columbia, Columbia, MO, 65212, USA; NextGen Precision Health Institute, University of Missouri-Columbia, Columbia, MO, 65212, USA; Department of Pediatrics, Washington University School of Medicine, Washington University in St. Louis, St. Louis, MO, USA

## Abstract

Recently, large-scale scRNA-seq datasets have been generated to understand the complex and poorly understood signaling mechanisms within microenvironment of Alzheimer’s Disease (AD), which are critical for identifying novel therapeutic targets and precision medicine. Though a set of targets have been identified, however, it remains a challenging to infer the core intra- and inter-multi-cell signaling communication networks using the scRNA-seq data, considering the complex and highly interactive background signaling network. Herein, we introduced a novel graph transformer model, PathFinder, to infer multi-cell intra- and inter-cellular signaling pathways and signaling communications among multi-cell types. Compared with existing models, the novel and unique design of PathFinder is based on the divide-and-conquer strategy, which divides the complex signaling networks into signaling paths, and then score and rank them using a novel graph transformer architecture to infer the intra- and inter-cell signaling communications. We evaluated PathFinder using scRNA-seq data of APOE4-genotype specific AD mice models and identified novel APOE4 altered intra- and inter-cell interaction networks among neurons, astrocytes, and microglia. PathFinder is a general signaling network inference model and can be applied to other omics data-driven signaling network inference.

## Introduction

Single-cell RNA sequencing data (scRNA-seq) technologies become popular in recent years because of their ability to profile gene expression and analyze cell composition in the single cell resolution^1–3^. On the one hand, by profiling and annotating scRNA-seq data, researchers can analyze differentially expressed genes in each cell population and sub-populations to understand which gene is altered in certain conditions. On the other hand, scRNA-seq data also shows great potential in discovering intra- and inter-cellular communication. However, there are only limited methods for discovering active signaling pathways or intra-cellular communication using scRNA-seq data. The existing models for doing this are mainly based on correlation, regression, and Bayesian analysis^4^, and the direct interaction signaling cascades usually were ignored in those methods because only a small set of genes have gene expression changes between different conditions^5^. For example, the CellPhoneDB^6^ can model the interactions between the ligands from one cell type to the receptors from another cell type. However, it cannot model the downstream signaling. CCCExplorer^7^ can discover both the ligand-receptor interaction and downstream signaling network by modeling differentially expressed genes. NicheNet^8^ takes a further step to integrate various interaction databases and train a predictive model to predict the interaction potential between the ligand and downstream targets. However, it only applies a statistical model, which cannot generate a clear communication path. CytoTalk^9^ applies the Steiner-tree to discover the de-novo signal transduction network from gene co-expression. However, the discovered signaling is based on co-expression and the physical interaction cascade is still unknown.

In the past few years, graph neural networks (GNNs) become famous due to their great performance on node and graph representation and classification tasks. For instance, GraphSAGE^10^ first proposed a general framework for learning the node representation in an inductive way. GAT^11^ incorporates the attention mechanism in GNN to actively learn how to aggregate all the information in graphs. The DGCNN^12^ model proposed sortPooling to efficiently sort nodes and learn graph features for graph classification. GIN^13^ proved that the normal message passing GNNs can only be at most as expressive as the 1-dimensional Wifelier-Lehman test (1-WL test) on learning graph structure and proposed a new GNN algorithm that is equally powerful as the 1-WL test. More recently, researchers have tried to generalize the transformer architecture^14^ into graph learning fields as it already shows superior power in learning both text and image data. Many works ^15–20^ show great potential for applying the transformer on the graph data. They either nest GNN architectures in the transformer layer, design specific attention mechanisms, or design novel encoding mechanisms to incorporate the graph structure into the transformer model. However, how to leverage GNNs to discover the intra- and inter-cell communication network is unknown as GNNs are typically a black-box model and it is hard to interpret the prediction results.

In this work, we present a novel framework called PathFinder to discover both intra- and inter-cell communication networks with a novel Graph transformer-based neural network. Given the scRNA-seq expression data and the condition (control/test), the PathFinder first samples a series of predefined paths through the prior gene-gene interaction database. Then, the PathFinder model takes the scRNA-seq expression data and the predefined path list as input to predict the condition of each cell. Through the training, the path important score will be learned to indicate the relative importance of each path in separating between the control and test conditions. To learn different types of communication like up-regulated or down-regulated networks, the novel regularization term is introduced to regularize the learned path scores to be close to the prior scores. After training. The path score will be sorted and the intra-communication network for each cell type will be generated by extracting the top K important paths. To generate the inter-cell communication network between the ligand cell and receptor cell, the intra-cell communication network will be collected for the receptor cell, and the ligand list will be extracted from the differential expressed gene list in the ligand cell. Finally, the ligands are linked to the intra-cell network with the ligand-receptor interaction database. The overall procedure of generating both intra- and inter-cell communication networks using PathFinder is shown in **Figure 1**. To our best knowledge, this is the first method to apply deep learning and graph transformers to discover signaling networks in scRNA-seq data. The advantages of PathFinder are listed below: (1) The model is designed based on a graph transformer, which has the great ability to learn both local- and long-range signaling patterns from gene expression and large-scale networks. (2) it is capable of identifying and providing the full signaling network between cells via cellular ligand and receptor. (3) The proposed PathFinder is a general framework, where users have the flexibility to input their own defined signaling paths or gene-gene interaction network database to identify important signaling based on user interest. Furthermore, (4) it can separate and generate different types of communication networks (Differential expressed/up-regulated/down-regulated), which allows more precise downstream analysis. We applied the PathFinder to a mice cohort of AD disease. The PathFinder not only achieves great prediction results but also generates intra- and inter-cell communication networks that align well with the latest knowledge on the mechanism of AD disease.

**Figure 1.**
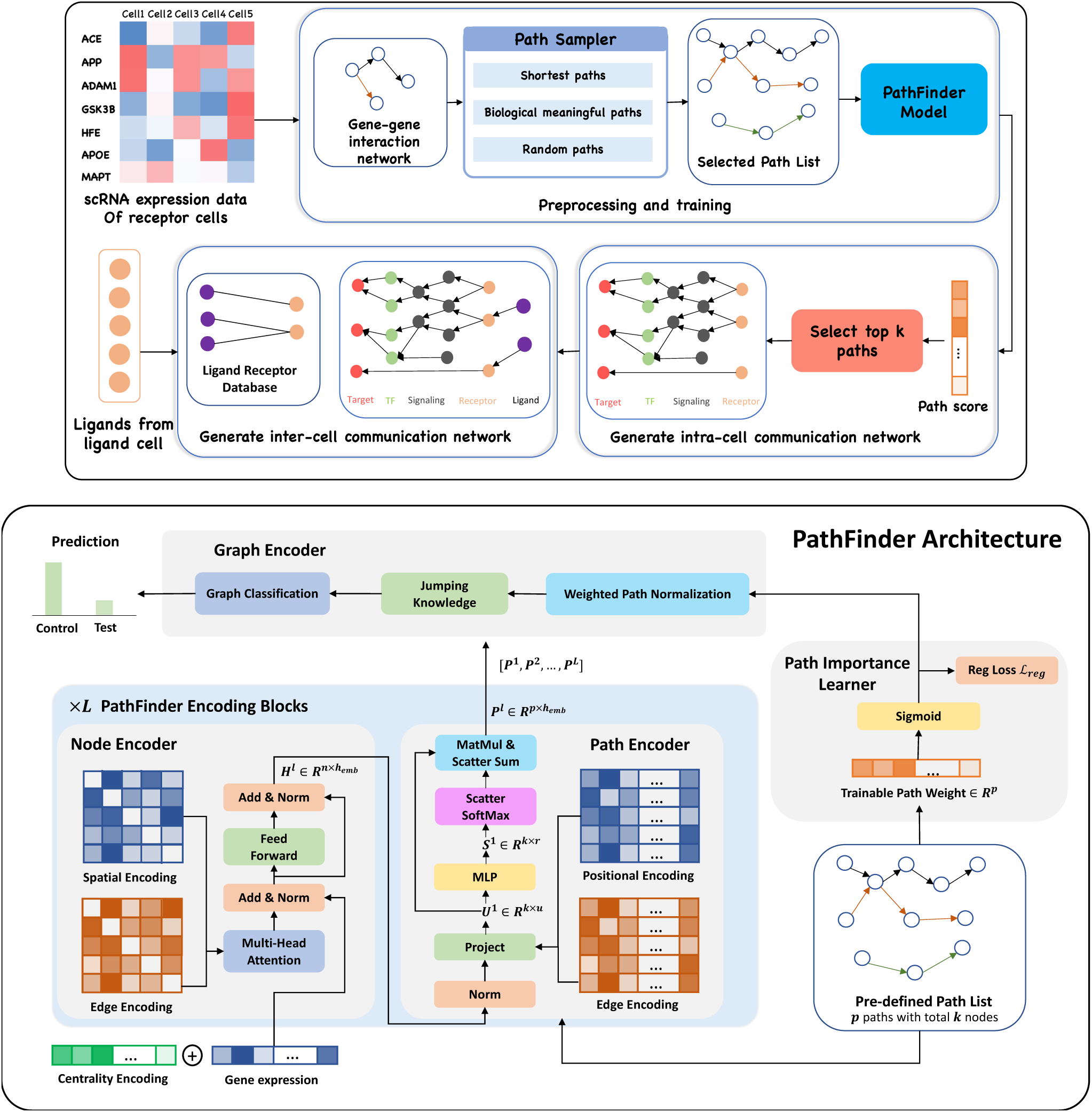
(Upper) Overview of PathFinder method to discover both intra- and inter-cell communication network. The input scRNA expression data with both samples from the control condition and the test condition is used to construct the gene-gene interaction network based on our large database. Next, the path sampler is used to generate all pre-defined path from the interaction network. Then, the PathFinder model is trained to separate the cells from two different conditions. After the training, the learned path score can indicate the importance of each path. Next, the top *k* paths are selected to generate the intra-cell communication network. Finally, the Ligand-receptor database is used to link all picked ligands (like differential expressed ligands) from ligand cells to the receptors in the intra-cell communication network of receptor cells to construct the inter-cell communication network. **(Lower) Model architecture of PathFinder.** The PathFinder consists of three components: node encoder, path encoder, and graph encoder. The node encoder is a stack of *L* layer of transformer with special encoding to encode local graph structure information of each node. The path encoder take the output from each layer of node encoder to learn long-range path embedding for each pre-defined path. Finally, the graph encoder aggregate information from each path to generate graph embedding and make final prediction. In the graph encoder, the trainable path weight will be learned to assign each path an importance score, which can be used to generate intra-cell communication networks.

## Results

### scRNA-seq data of Alzheimer’s disease cohort on mice

To evaluate the proposed PathFinder method, scRNA-seq data of Alzheimer’s disease is collected from the Gene Expression Omnibus (GEO) database with accession number GSE164507^21^. The raw data is processed using the Seurat R package^22^ and the process procedure is done by following the previous study^21^. Specifically, we select cell samples from two different conditions, denoted as TAFE4_tam and TAFE4_oil. TAFE4_tam stands for mice with the APOE4 gene be knocked out from astrocyte cells and TAFE4_oil stands for the mice with the existence of APOE4. It is well known that APOE4 is one of the most significant genetic risk factors for late-onset AD. However, the detailed mechanisms behind the APOE4 are still unknown. By analyzing the difference between the signaling pattern with and without APOE4, we can get a deeper understanding of the effects of the APOE4 gene on brain cells. Concretely, the excitatory neuron (Ex), microglia (Mic), and astrocyte (Ast) of the TAFE4 group are collected from the dataset with a total number of samples of 13604, 3874, and 734, respectively. Then, the PathFinder method is applied to predict the condition of each cell (oil or tam) separately for each cell type and generate both intra- and inter-cell communication networks between these three cell types. The pre-defined path list including all the shortest paths starting from receptors and all possible paths from the receptor to the target gene. For the shortest distance paths, we only select paths with a minimum length of 3 (except all receptor direct regularizations, which has a length of 2) and a maximum length of 10. We compute the prior weight of each path based on the average differential expression level of all genes in the path (more details in the Method section) for the path score regularization. To ensure the robustness of the analysis, we only selected the top 8192 variable genes from the original dataset as input to the model, which resulted in the final number of pre-selected paths being 1210.

### PathFinder can effectively separate cells from different conditions by selecting differentially expressed signaling paths

To evaluate the performance of the PathFinder model, the model is applied to excitatory neurons, astrocytes, and microglia cells separately to predict the conditions of each cell (tam/oil), denoted as TAFE4_ex, TAFE4_mic, and TAFE4_ast respectively. For each cell type, we repeat the training 5 times with each time splitting the dataset into train, validation, and test subsets randomly with a ratio of 0.7/0.1/0.2. We report the average performance and standard deviation on the test set over all 5 runs. The detailed experimental setting can be found in the Method section. The detailed results are shown in **Table 1** and **Figure 2a**.

**Figure 2.**
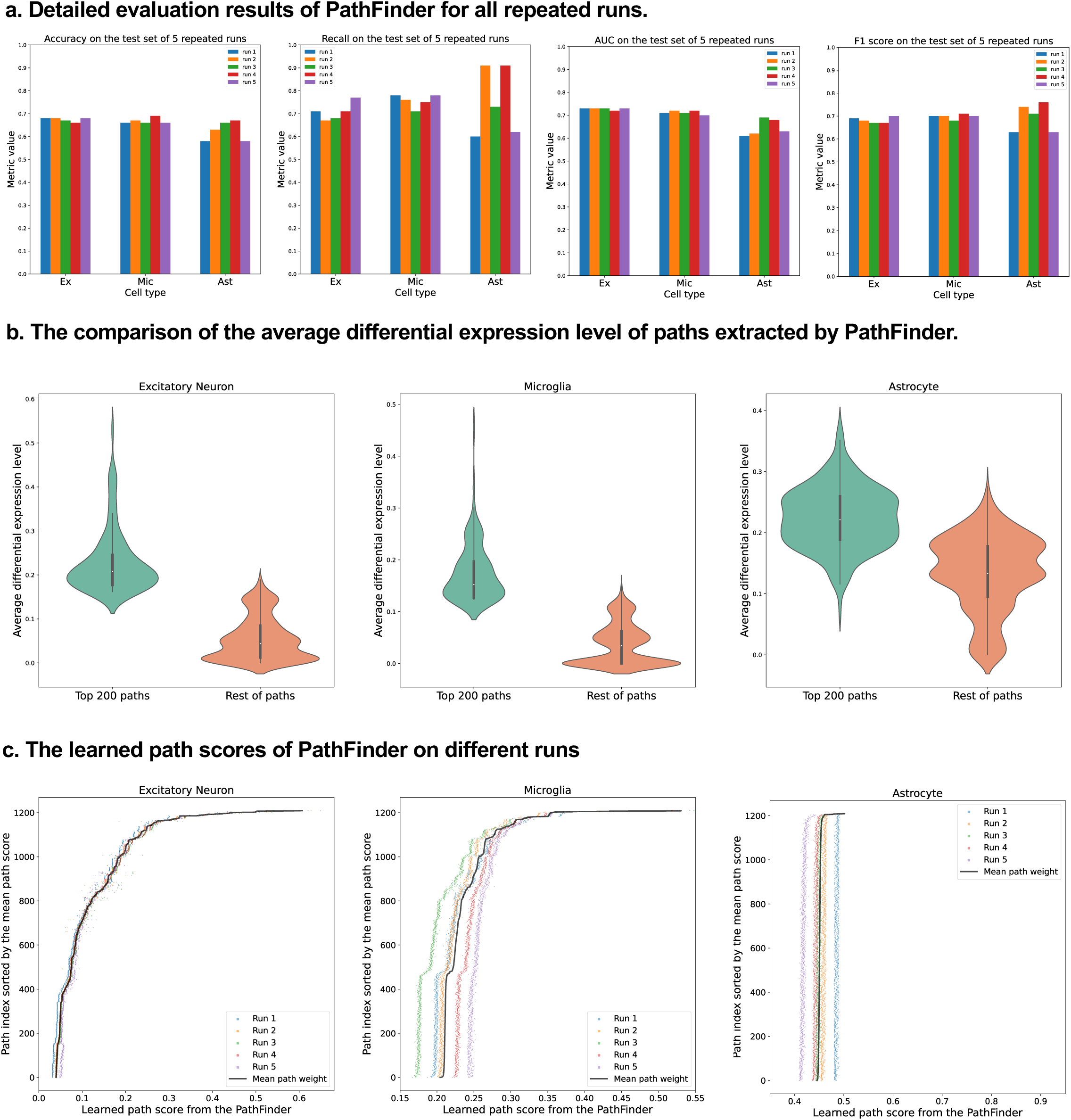
Evaluation of the PathFinder model. **(a)** The detailed evaluation metrics on test dataset from all runs. **(b)** The comparison of the average differential expression level of top paths sorted by PathFinder during the training. The top 200 paths have higher differential expression level than others for all three cell types. **(c)** The learned path scores of PathFinder on different runs. All paths are ranked by the average score across all runs.

**Table 1:** Evaluation results of the PathFinder model.

|  | **Accuracy** | **Recall** | **Precision** | **Specificity** | **F1** | **AUC** |
| --- | --- | --- | --- | --- | --- | --- |
| **TAFE4_ex** | 0.67±0.01 | 0.71±0.04 | 0.66±0.02 | 0.64±0.05 | 0.68±0.01 | 0.73±0.01 |
| **TAFE4_mic** | 0.67±0.01 | 0.76±0.03 | 0.65±0.02 | 0.58±0.04 | 0.70±0.01 | 0.71±0.01 |
| **TAFE4_ast** | 0.62±0.04 | 0.75±0.15 | 0.65±0.03 | 0.44±0.14 | 0.69±0.06 | 0.65±0.04 |

As can be seen, the PathFinder can successfully classify the majority of cells in the test dataset into the correct condition. This means that after training, the model learned the most important difference between the two conditions from a huge gene expression profile. Such differences can be reflected in the important score of each path as the final prediction is made based on the different predefined paths. Among all results, the standard deviation of the metrics for TAFE4_ast is much larger than the other two cell types. We conjecture this is caused by the limited number of cell samples in the TAFE4_ast group, which makes the model easily overfit to the training data. Next, we evaluate the learned path score from each group. For each cell group, we first average the learned path score from 5 repeated runs to get the final path score. Next, we average the absolute fold-change level of all genes within each path to get an average differential expression level for each path. Then, we compare the top 200 selected paths from the results of PathFinder model to the rest of the paths. The results are shown in **Figure 2b**. We can see that for all three different cell types, the selected top 200 paths from PathFinder have much higher average differential expression level comparing to the rest of paths. The results indicate that PathFinder is effective for ranking differential expressed paths through the training. This can be attributed to two objective function used in the PathFinder. First, by minimizing the classification loss, the model are forced to increase the score for paths that are useful for separating two different conditions. It is intuitive that paths with higher average differential expression level are more helpful for the prediction. Second, by minimizing the regularization loss, the model tends to give a high score for paths with high prior weight, and the prior weight is positively related to the average differential expression level.

Then, we evaluate the robustness and stability of the PathFinder. Concretely, we want the final path score distribution (ranking) learned from PathFinder to be stable and robust even if we slightly alter the training data. Since we randomly split the whole dataset for each repeated run, we can directly compare the learned score for each run to achieve our goal. Therefore, we plot the learned score for all paths and all runs with paths are sorted by the average score. The results are shown in **Figure 2c**. We can see for all three cell types, the learned scores are very stable across different runs, as paths with higher ranks always have higher scores. This means that even if we slightly alter the training dataset, PathFinder model can still output almost the same top k paths. The results successfully demonstrate the robustness of the PathFinder for extracting important paths and constructing intra-communication networks.

### Core intra-cell signaling networks associated with APOE4 genotype

In this section, we evaluate the intra-cell communication networks discovered by PathFinder model. Particularly, we want to know whether the discovered networks can reveal the recent discovery of APOE4-driven AD or even indicate new findings. First, for all three cell types, the final networks are generated by first averaging the path score learned from 5 repeated runs and then ranking and selecting the top 300 paths from all paths to form the final networks. The generated networks for all three cell types are shown in **Figure 3**. Next, we perform the enrichment analysis on all generated networks using KEGG signaling pathways and Gene Ontology (GO) terms. The enrichment results are shown in **Figure 4a**. Based on the results, we find several key factors that are important to the development of APOE4-driven AD.

**Figure 3.**
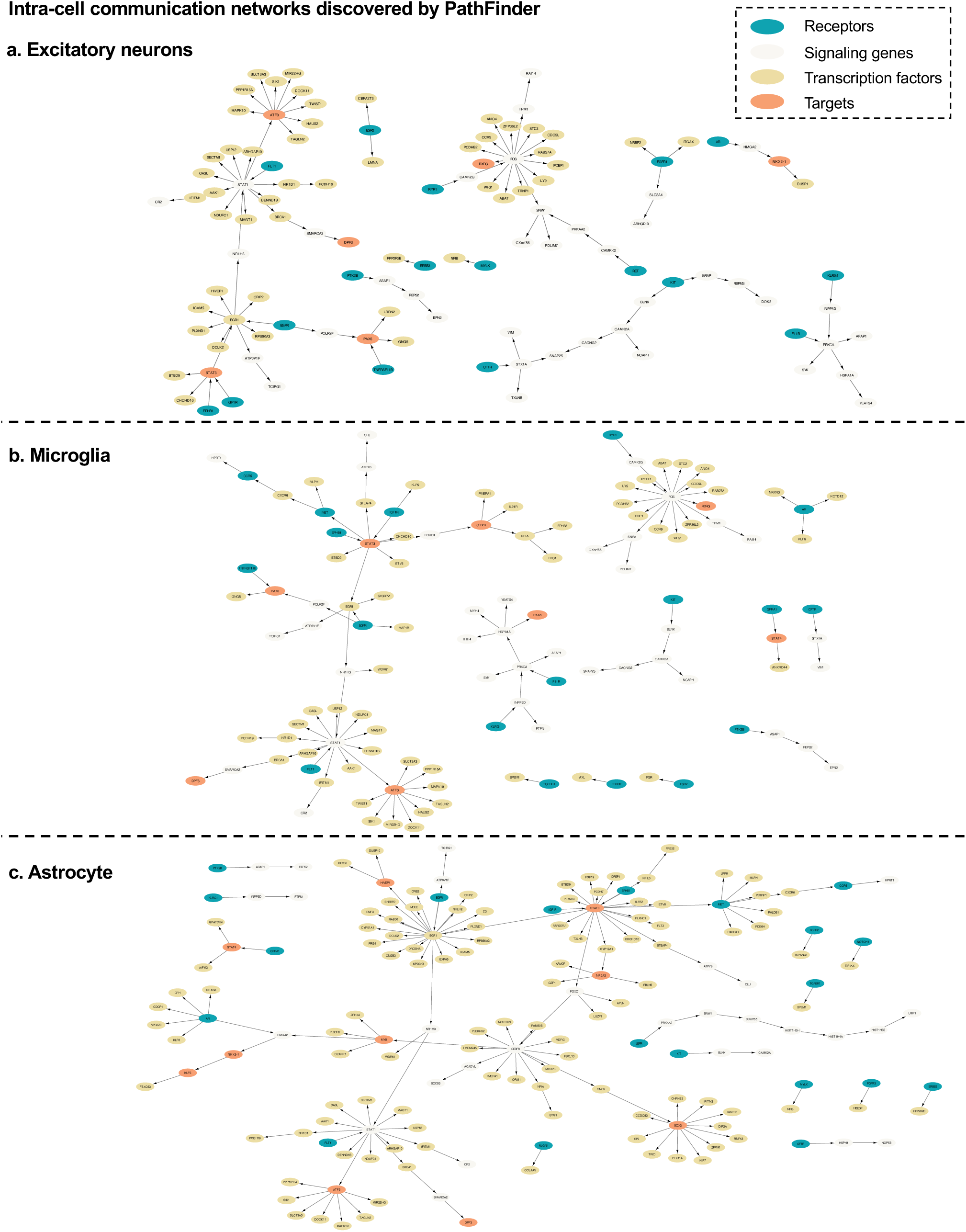
Intra-cell communication networks discovered by PathFinder model.

**Figure 4.**
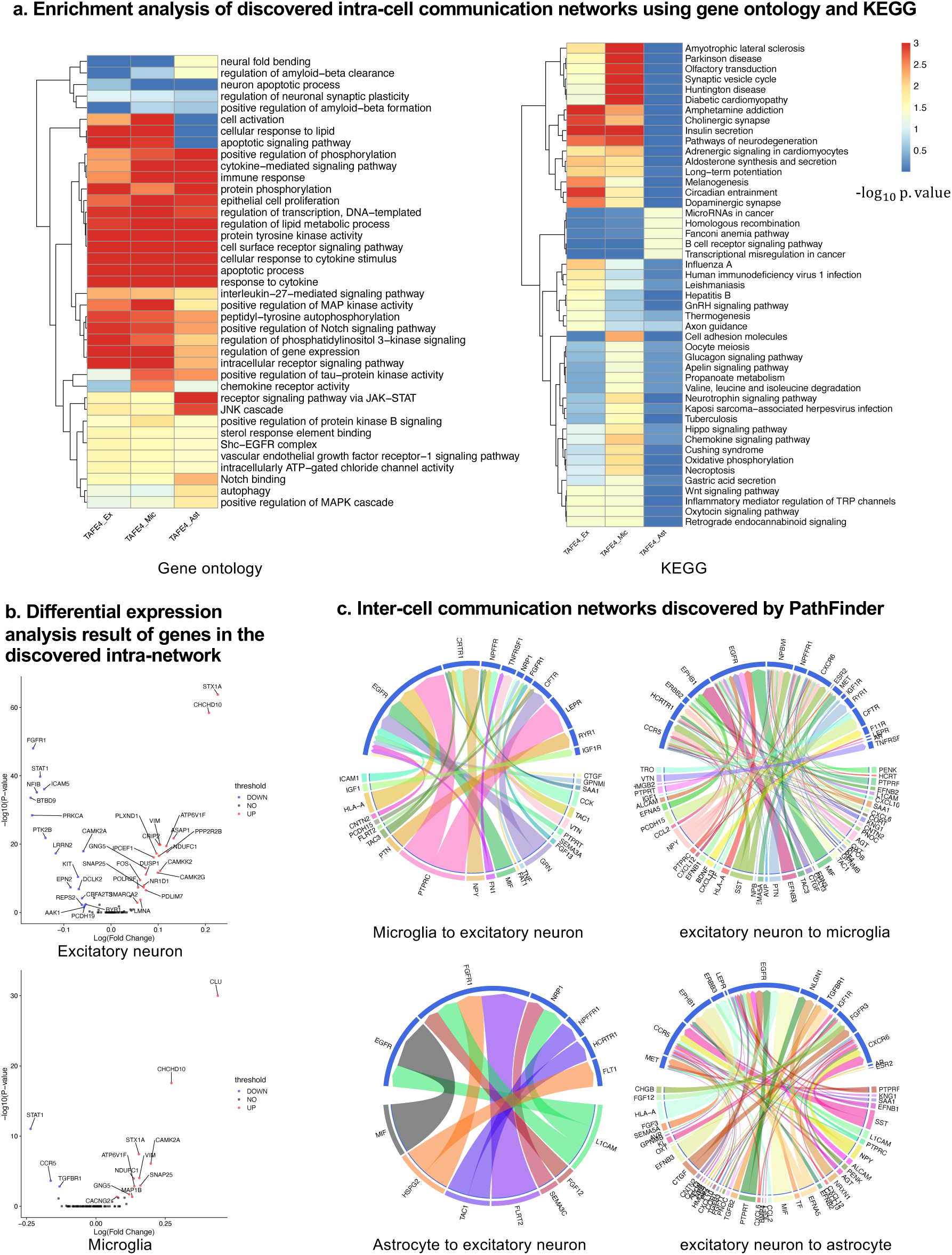
Analyses of the results. **(a)** KEGG and GO enrichment analysis on all discovered intra-networks. **(b)** Differential expression analysis. **(c)** Inter-cell communication networks. All ligands are from DEGs of the ligand cells. Receptors are marked as blue.

#### Neuron inflammation

Numerous studies have shown that inflammation is highly activated and plays a key role in the progress of AD^23–26^. From the enrichment results, we can see that many inflammation-related pathways/GO terms are enriched across multiple cell types. For example, *cytokine-mediated signaling pathway, cellular response to cytokine stimulus, inflammatory mediator regulation of TRP channels.* This result aligns with the previous studies and further confirms that the existence of APOE4 in the astrocyte stimulates the inflammatory response. More specifically, several genes related to neuron inflammation are identified by PathFinder across multiple cell types. STAT1 and STAT3 are identified as hub genes connected to multiple targets in both the network of neurons and microglia. It has shown that STAT1 plays a key role in the regulation of inflammatory response and cellular death^27,28^. Moreover, the differential expression analysis (**Figure 4b**) reveals that STAT1 is highly differential expressed in TAFE4 group, which further confirms the important role of STAT1.

#### Autophagy

Besides the inflammation, the *Apoptotic* and *Apoptotic signaling pathway* are enriched in the neuron and the Microglia. The autophagy is a lysosome-dependent, homeostatic process, in which organelles and proteins are degraded and recycled into energy. Autophagy has been linked to Alzheimer’s disease pathogenesis through its merger with the endosomal-lysosomal system, which has been shown to play a role in the formation of the latter amyloid-β plaques^29^. One hypothesis states that irregular autophagy stimulation results in increased amyloid-β production^30^. The existence of the APOE4 may also affect the process of autophagy and thus cause the acclamation of the amyloid-β of the AD’s brain. Particularly, genes CLU and FOXO1 are identified in the intra-network of microglia. CLU is one of the top AD candidate genes. Some study shows that it is a casual gene of AD-affected hippocampal connectivity^31^. Moreover, it is shown that CLU protein interacts with Aβ, reduces its aggregation, and protects against its toxic effects^32^. Many studies have shown that FOXO1 induces autophagy in cardiomyocytes and cancer cells. FOXO1 was also reported to be a gene encodes transcription factor that plays a role in autophagy modulation in neurons^33^.

#### Lipid transportation

*The regulation of lipid metabolic process and cellular response to lipid* are enriched in the intra-communication network of all three cell types. The enriched gene included: NR1D1, EGR1, BRCA1. It has been proved that the APOE4 is involved in the lipid transport and metabolism^34^. The existence of the APOE4 in the astrocyte may disturb the brain lipid composition and thus affect the blood-brain barrier (BBB) function^35^. All these results confirm the influence of the APOE4 in the progress of AD and the dysfunction and death of the neuron.

#### JAK-STAT signaling pathway

In the intra-communication network of the astrocyte, the receptor signaling pathway via JAK-STAT is enriched with the corresponding gene: STAT3, SOCS3, HMGA2, STAT1. The JAK-STAT signaling pathway has been reported to be the inducer of astrocyte reactivity^36^. The enrichment of the pathway indicates that the existence of the APOE4 in astrocytes can influent the function of the JAK-STAT signaling pathway and the pathway reversely affects the activity of the astrocyte.

### Core multi-cell inter-cell communication networks associated with APOE4 genotype

To further understand the complex signaling flow and mechanism behind the APOE4 and AD pathology, we further generate inter-cell communication networks between three different cell types using the PathFinder, as shown in **Figure 4c**. First, we can see that compared to astrocytes, microglia have much more interactions with neurons. This may indicate that the existence of APOE4 in the astrocyte may activate the functionality of microglia and then cause abnormal activities in the neurons.

Among all interactions, several interesting interactions appealed to the result. First, the MIF secreted by the astrocyte interacts with the EGFR in the neuron and follows downstream signaling. The MIF is a well-known proinflammatory cytokine that promotes the production of other immune mediators. Increased expression of MIF can contribute to chronic neuroinflammation and neurodegeneration^37^. EGFR has been shown to be a potential target for treating AD-induced memory loss^38,39^. The increased expression level of MIF could be the signature of activated astrocytes and the MIF further triggers the expression of EGFR and following downstream network in the neuron, which contributes to neuron inflammation and degeneration.

Besides MIF in astrocyte, many ligands for EGFR receptor are also identified in Microglia, including ICAM1, IGF1, HLA-A, CNTN2, PCDH15, FLRT2, TAC3, PTN, and PTPRC. The down-regulation of PTPRC is reported to be contributed to the overproducing of A β and neuron loss^40^. Another interaction is the *NLGN1* which is expressed in neurons interact with the *NRXN1* gene in the astrocyte. The amyloid-β oligomers are synaptotoxins that build up in the brains of patients and are thought to contribute to memory impairment in AD. It has been shown that the interaction neurexins (Nrxs) and neuroligins (NLs) is critical for synapse structure, stability, and function^41^. The dysregulation of the interaction between Nrxs and NLs may contribute to the formation of amyloid-β oligomer. The *EFNA5* in the neuron is up-regulated in the neuron and interacts with the *EPHB1* and downstream *STAT3* signaling in the astrocyte. This interaction is closely related to the ephrin-B1-mediated stimulation. Analysis has shown that the ephrin-B1-mediated stimulation induces a protective and anti-inflammatory signature in astrocytes and can be regarded as “help-me” signal of neurons that failed in early amyotrophic lateral sclerosis (ALS)^42^. Such signals could also play an important role in triggering inflammation and neuron degeneration in CNS system.

### Conclusions and Discussions

In this work, we propose PathFinder, which is the first deep-learning model with a graph transformer that can be used to extract both intra- and inter-cell communication network with scRNA-seq data. Through a case study using an AD scRNA-seq dataset from mice, we evaluate the effectiveness of PathFinder from multiple perspectives. First, the quantitative analysis confirm that PathFinder obtains great performance in separating cells from different conditions by leveraging the difference of expression patterns in the signaling paths. Further, the learned path score is robust and consistent in repeat runs. We further evaluate the correctness of extracted networks through extensive literature searches. The resulting network aligned well with many recent discoveries on AD pathology, which further proved the effectiveness of the proposed PathFinder. Also, the current version of PathFinder has a few potential limitations to be improved in the future work. First, it requires many samples in training to produce reasonable results. Second, it relies on the pre-defined paths from the database to learn and extract meaningful patterns and is unable to discover new signaling flows. All these limitations are worth to further investigate, and we will improve the model in our future work.

## Methodology

### Gene-gene interaction database collection and processing

To construct the gene-gene interaction database, the raw interaction data was collected from NicheNet software.^8^ The raw interaction data were divided into three types: ligand-receptor network, signaling network, and gene-regulation network. The original network contains 12019 interactions/1430 genes, 12780 interactions/8278 genes, and 11231 interactions/8450 genes, respectively. To construct the intra- and inter-network database, the data was further processed by the following steps.

First, ligands and receptors were collected by gathering the source and target of the ligand-receptor network. There are a total of 688 ligands and 857 receptors. Then, interactions in the ligand-receptor network were divided into two types. If one interaction exists in both directions in the database, we labeled it as bidirectional. Otherwise, we labeled it as directional. After processing, there are 11880 directional interactions and 139 bidirectional interactions.

The gene-regulation network was processed as follows. First, 1639 transcriptional factors (TFs) were collected from51. For convenience, TFs that exist in either ligand or receptor list were removed. Finally, 1632 TFs were collected. Next, three different types of regulation were collected in the gene-regulation interaction network, which are ligand regulation, receptor regulation, and TF regulation. To label each interaction into one of three types, all the interactions in the network were removed if the source gene was not in either ligand, receptor, or TF list. Then, the interactions were labeled based on the type of source (e.g. if the source of interaction is a receptor, we label it as receptor regulation). After processing, there are 1329 ligand-regulation interactions, 272 receptor-regulation interactions, and 6706 TF-regulation interactions.

Finally, the signaling network was processed as follows. First, all the interactions were removed if they existed in either the ligand-receptor or the gene-regulation network. Then, the interactions were divided into receptor-TF, receptor-signaling, signaling-TF, and signaling-signaling further. To be more specific, if the source of interaction is in receptor list and target of interaction is in TF list, the interaction was labeled as receptor-TF. If the source of interaction is in receptor list and target is not in TF list, the interaction was labeled as receptor-signaling. If the source of interaction is not in receptor list and target of interaction in TF list, the interaction was labeled as signaling-TF. If neither the source nor target of interaction are in the TF and receptor list, the interaction was labeled as signaling-signaling. The interactions cannot be classified into one of above group will be removed for connivence. Finally, there are 31 receptor-TF interactions, 524 receptor-signaling interactions, 975 signaling-TF interactions, and 9745 signaling-signaling interactions.

### Notations

A gene graph is denoted as *G* = (*V*, *E*), where *V* is the set of gene nodes with |*V*| = *n*, *E* is the set of edges and *E* ⊆ *V* × *V*. The node feature set is denoted by *X* = [*x*_1_, *x*_2_, …, *x*_*n*_]^*T*^ ∈ *R*^*n*×*d*^, where *x*_*u*_ ∈ *R*^*d*^ is the feature vector of the node *u*. The graph structure is defined by an adjacency matrix *A* ∈ [0,1]^*n*×*n*^, where *A*_*uv*_ = 1indicate there is an edge from the node *u* to node *v* and *A*_*uv*_ = 0 otherwise. Further, a set of paths sampled from graph is denoted as *P* = {*p*_1_, *p*_2_, …, *p*_*p*_}, where *p*_*m*_ is the *m*-th path, which is a list to store the nodes of the path in order. Paths can have different lengths, and we denote the length of path *m* be *l*_*m*_.

### Preliminary of transformer and Graphormer

The transformer is a powerful architecture in the deep learning field. It consists of multiple transformer layers. Each transformer layer has two parts: a multi-head self-attention and a point-wise feed-forward network (FFN) with residual connection applied between each part. Let 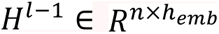 be the embedding of nodes in layer *l* − 1, and 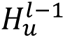 is the embedding of the node *u* in layer *l* − 1, the computation of multi-head self-attention is :

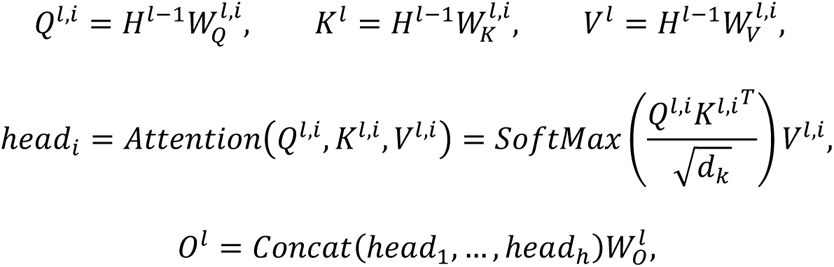

where 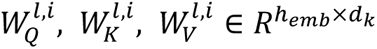, and 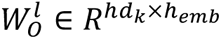 are all trainable weight matrix, *h* is the number of heads, 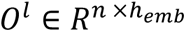 is the output from the multi-head self-attention in layer *l*. For simplicity, we let *h* × *d*_*k*_ = *h*_*emb*_. The output *O*^*l*^ will then be fed into a point-wise feed-forward network. The computation of the point-wise feed-forward network is:

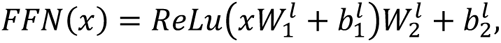

where 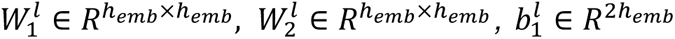, and 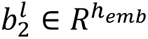 are all trainable weight matrix and bias. Notice that here we slightly modify the hidden size of the feed-forward network of the original model.

However, the vanilla transformer cannot be used directly on the graph structure data as it lacks critical part for encoding the topological information into model. To deal with this issue, Graphormer proposed several novel encodings into the model. Specifically, they introduced centrality encoding, spatial encoding and edge encoding. The centrality encoding is used to embed the graph centrality information into the model. Given the input data *X*, the computation of centrality encoding is:

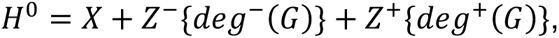

where the *Z*^−^, *Z*^+^ are all trainable embedding vectors and *deg*^−^(*G*), *deg*^+^(*G*): *G* → *R*^*n*^ are the function to compute the in-degree and out-degree of each node in the graph *G*. The spatial and edge encoding is used to encode the graph structure into the model. With the spatial and edge encoding, the self-attention is revised as:

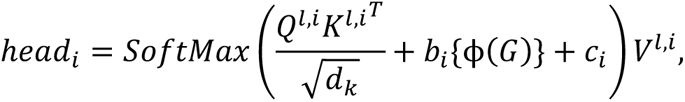

where *b*^*i*^ is trainable embedding vectors to encode the spatial information at head *i* and *ϕ*(*G*): *G* → *R*^*n*×*n*^ is the function to compute the shortest path length between each two nodes. If two nodes are not connected, a special value will be used. *c*^*i*^ ∈ *R*^*n*×*n*^ is the edge embedding and 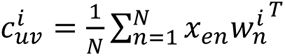, where *x_en_* is the edge feature of the *n*-th edge in the shortest path between node *u* and node *v* and the 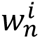 is trainable weight vector of *n*-th edge of head *i*. Note that both the spatial and edge encoding are unique across different layers.

### Architecture of the PathFinder

The PathFinder model consists of three components. Namely, the node encoder, path encoder, and graph encoder. The overall architecture of the PathFinder model is shown in **Figure 1-lower.** The architecture of the node encoder is similar to the Graphormer, which stacks *L* transformer layer with centrality encoding, spatial encoding and edge encoding. The input to the PathFinder is the expression value of each gene in a cell sample. However, we made several modifications to the original architectures. First, the hidden size in the point-wise feed forward network is all *h*_*emb*_ in both two layers for simplicity. Second, the edge encoding in the PathFinder is modified. In the original Graphormer, the edge encoding is computed by all the edges in the shortest path between two nodes, which can capture long-range information in the graph. However, the localized feature in the graph will be smoothed in such a manner. Instead, the PathFinder want the node embedding learned from the node encoder to focus on the localized information in the graph. Therefore, the direct edge encoding is proposed. The direct edge encoding is computed by:

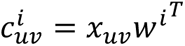

Where *x*_*uv*_is the edge feature of the edge between node *u* and node *v*. If there isn’t an edge between two nodes, the direct edge encoding is set to a special vector for simplicity. By doing this, the node encoder is good at learning node embedding with localized information. Finally, the spatial encoding is also revised in the PathFinder. Since here the graph structure is identical for all samples and the node order invariant is automatically held, we can learn a specific spatial encoding for each pair of two nodes. Therefore, we design the node index encoding in the PathFinder. The node index encoding is not computed from the length of the shortest path between each pair of nodes but is directly learned for each pair of two genes. Namely, for each pair of two genes, a unique encoding is learned for each head in each layer of the node encoder. Next, the path encoder is responsible for learning gene signaling path embedding. Given the node embedding in the graph and the pre-defined path list of the graph. The details of the pre-defined path list are illustrated below. Suppose there are *p* unique paths in the path list *P*, where the length of the *m*-th path is *l*_*m*_ and the total number of nodes in the path list is *k* (count repeated nodes in different paths). Denote the node embedding output from the layer *l* as *H*^*l*^, we first learn a path-specific embedding through:

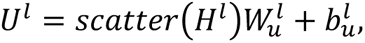

where 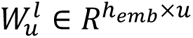 and 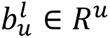 are all trainable weight matrix, *scatter* is a function to reorder and scatter the node in the graph into the order of the pre-defined path list. *U*^*l*^ ∈ *R*^*k*×*u*^ is the learned path-specific embedding. Then, the positional encoding and edge encoding are introduced to encode additional information for all paths. Let *U*^*l*^ be the result embedding after the special encodings. We have:

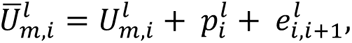

Where 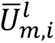 and 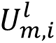 are the embedding of *i* -th node in the *m* -th path, 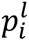 is the learnable positional encoding vector and its value only depends on the position *i*, 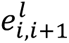 is the learnable edge encoding to encode the edge type between *i*-th node and *i* + 1-th node. Next, the score of each node within the path is computed by:

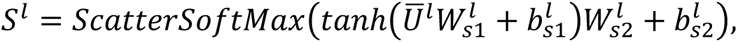

where the 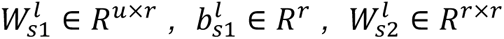, and 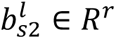 are all trainable parameters. *ScatterSoftmax* is the softmax function working on each path and each dimension. The *S*^*l*^ ∈ *R*^*k*×*r*^is the final *r* set important score for each node in each path. We let the *r* × *u* = *h*_*emb*_ for simplicity. After we obtain the *S*^*l*^, the path embedding is computed by:

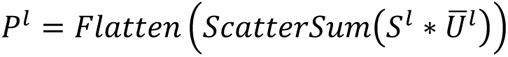

The ∗ is the point-wise product working on each set of important scores. That is, for each set of important score, we do a point-wise product of that set of score and *U*^*l*^, which result in total *r* sets. The *ScatterSum* function is the summation on each path. The *Flatten* is the function to flatten the embedding of all sets. The 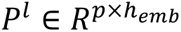 is the final path embedding in the layer *l*.

In the original Graphormer, the graph embedding is learned by introducing a special node and letting it connect to all the nodes in the graph. After forwarding, the embedding of that special node is regarded as the graph embedding for the graph-level task. In PathFinder, we want to learn the graph embedding from the path embedding. Meanwhile, we want to extract the important paths from the model after we train the model given the graph-level task. To simultaneously achieve both two goals, the graph encoder is proposed. The graph encoder consists of two parts, the first part is a trainable path weight and the sigmoid function to assign each path with different scores. The second part is the jumping knowledge network to combine the graph embedding in each layer and compute the final embedding.

In PathFinder, the graph embedding is learned by integrating all the path embedding from each layer, which requires an important score for each path. Normally, the score is computed based on one sample. However, such a score is not robust and may vary a lot even given a minor variation of the path embedding^44,45^. To avoid the issue and learn a robust important score across the whole dataset, the trainable path score *M* ∈ *R*^*p*^ is introduced. *M* is identical to all samples and layers and learned through back propagation. The path important score is computed by:

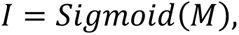

where *I* ∈ *R*^*p*^ is the important score for each path. Next, the graph embedding of layer *l* is computed by:

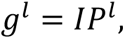

where *g*^l^ is the graph embedding of layer *l*. The final step of the graph encoder is to integrate the graph embedding of each layer and learn a final embedding. Here we utilize the idea of JumpingKnowledge network^46^ and compute the final graph embedding by:

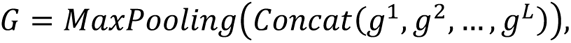

where *MaxPooling* is the max pooling function and 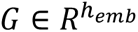 is the final graph embedding learned by the PathFinder. Finally, the graph embedding is used to classify the cell sample into the corresponding condition (control/test). The prediction is a typical binary prediction compute by:

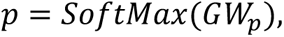

Where 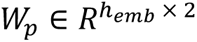 is the trainable projection matrix and *p* is the predicted distribution.

### Training and Regularization of the PathFinder

To train the PathFinder model, the negative log likelihood (NLL) loss is applied. Let the 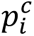 be the predicted probability of the true condition of cell *i*, then the NLL loss is computed by:

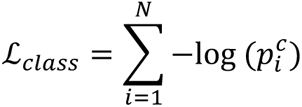

Where the *N* is the number of cells in the dataset. Meanwhile, to regularize the training of model and learn biological meaningful paths from the model, the regularization term is introduced to the path score *M*. Intuitively, the path that have higher total fold change should have higher path score. Further, we designed three different regularization terms to generate different important paths by introduce the prior path score. Specifically, these three regularizations are up-regulated path, down-regulated path, and differential expressed path regularization. Let the 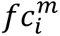 be the log fold change of gene *j* in path *m*, then the prior path score is computed by:

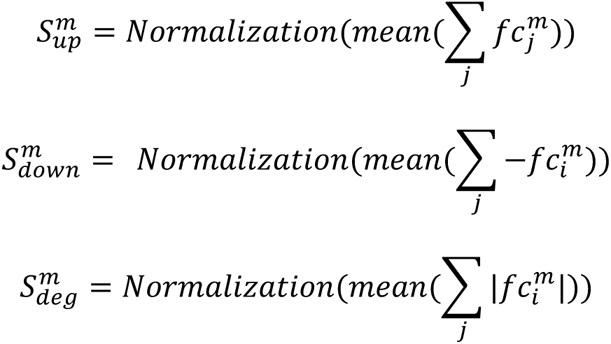

Where the 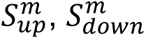, and 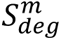 are the prior path score for up-regulated, down-regulated and differential expressed regularization respectively, *Normalization* is the min-max normalization across all paths. Suppose we use the up-regulated prior score, the regularization loss is computed by:

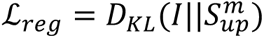

The final loss is:

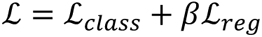

Where *β* is the weight of the regularization term.

### Predefined path list

To train the PathFinder, the path list needs to be defined before the training. Given the collected gene-gene interaction database, we designed several choices to generate a predefined path list. The first choice is the shortest path. For this choice, the shortest path between each pair of genes in the dataset will be computed and collected given the gene-gene interaction network. The second choice is to generate all the possible paths that start from the receptor and end in the target, which can also be done using the gene-gene interaction database. To constrain the path, the minimum length of the path is set to be 3 unless the path is a receptor regulation interaction. The maximum length of the path is set to be 10.

### Experimental details

We conduct experiments to validate the effectiveness of PathFinder on TAFE_ex, TAFE_mic, and TAFE_ast cell sample datasets. For each dataset, we randomly split datasets into train/validation/test sets with a ratio of 0.7/0.1/0.2. We train the model using the train set and validate the performance of the model using the validation set. Finally, we save the model that achieves the best performance on the validation set and report the performance of the saved model on the test set. We use the Area Under the Curve (AUC) as the performance metric for selecting the best model. We repeat experiments on each dataset five times (with each time a different random split on the dataset) and report the mean results and the standard deviation. The model and training hyperparameters are described as follows: we set the number of layers as 6, the hidden size *h*_*emb*_ as 128, the number of head, and the number of scores set *r* as 8. For each experiment, we set the number of training epochs as 30, the learning rate as 0.0005, the dropout rate as 0.1, the regularization weight *β* as 0.1 for TAFE_ex and TAFE_mic, and 1.0 for TAFE_ast.

### Generation of the intra- and inter-cell communication network

After the PathFinder model is trained, the generation of an intra-cell communication network is straightforward. Concretely, we first average the path weight learned from 5 repeated experiments to get the final path weights. Next, the top *K* paths are extracted and combined to generate the intra-cell communication network. The generation of the inter-cell communication network is as follows. Let the cell that provides ligands be the ligand cell and the cell providing receptors be the receptor cell. The intra-cell communication network is first generated. Then, the ligands of the ligand cell and receptors of the receptor cell will be extracted from the intra-network respectively. Then, the ligand-receptor pairs are selected given the ligand-receptor database. Finally, the kept pairs will be linked and the inter-network is generated.

## Acknowledgment

This study was partially supported by NIA R56AG065352 (to Li), 1R21AG078799-01A1 (to Li).

## Notes

### Competing Interest Statement

The authors have declared no competing interest.

